# The Rosetta all-atom energy function for macromolecular modeling and design

**DOI:** 10.1101/106054

**Authors:** Rebecca F. Alford, Andrew Leaver-Fay, Jeliazko R. Jeliazkov, Matthew J. O'Meara, Frank P. DiMaio, Hahnbeom Park, Maxim V. Shapovalov, P. Douglas Renfrew, Vikram K. Mulligan, Kalli Kappel, Jason W. Labonte, Michael S. Pacella, Richard Bonneau, Philip Bradley, Roland L. Dunbrack, Rhiju Das, David Baker, Brian Kuhlman, Tanja Kortemme, Jeffrey J. Gray

## Abstract

Over the past decade, the Rosetta biomolecular modeling suite has informed diverse biological questions and engineering challenges ranging from interpretation of low-resolution structural data to design of nanomaterials, protein therapeutics, and vaccines. Central to Rosetta’s success is the energy function: amodel parameterized from small molecule and X-ray crystal structure data used to approximate the energy associated with each biomolecule conformation. This paper describes the mathematical models and physical concepts that underlie the latest Rosetta energy function, *beta_nov15*. Applying these concepts,we explain how to use Rosetta energies to identify and analyze the features of biomolecular models.Finally, we discuss the latest advances in the energy function that extend capabilities from soluble proteins to also include membrane proteins, peptides containing non-canonical amino acids, carbohydrates, nucleic acids, and other macromolecules.

## Introduction

Proteins adopt diverse three-dimensional conformations to carry out the complex mechanisms of life. Their structures are constrained by the underlying amino acid sequence^1^ and stabilized by a fine balance between enthalphic and entropic contributions to non-covalent interactions.^2^ Energy functions that seek to approximate the energy of these interactions are fundamental to computational modeling of biomolecular structures. The goal of this paper is to describe the energy calculations used by the Rosetta macromolecular modeling program:^3^ we explain the underlying physical concepts, mathematical models, latest advances, and application to biomolecular simulations.

Energy functions are based on Anfinsen’s hypothesis that native-like protein conformations represent unique, low-energy, thermodynamically stable conformations.^4^ These folded states reside in minima on the energy landscape, and they have a net favorable change in Gibbs free energy, which is the sum of contributions from both enthalpy (∆*H*) and entropy (∆*S*) relative to the unfolded state. To follow these heuristics, macromolecular modeling programs require a mathematical function that can discriminate between the unfolded, folded, and native-like conformations. Typically, these functions are a linear combination of terms that compute energies as a function of various degrees of freedom.

The earliest macromolecular energy functions combined a Lennard-Jones potential for van der Waals interactions^5–67^ with harmonic torsional potentials^8^ that were parameterized using force constants from vibrational spectra of small molecules.^9–11^ These formulations were first applied to investigating the structures of hemolysin,^12^ trypsin inhibitor, ^13^ and hemoglobin^14^ and have now diversified into a large family of commonly used energy functions such as AMBER,^15^ DREIDING,^16^ OPLS, ^17^ and CHARMM.^18,19^ Many of these energy functions also rely on new terms and parameterizations. For example, faster computers have enabled the derivation of parameters from *ab initio* quantum calculations.^20^ The maturation of X-ray crystallography and NMR protein structure determination methods has enabled development of statistical potentials derived from per-residue, inter-residue, secondary-structure, and whole structure features.^21–28^ Additionally, there are alternate models of electrostatics and solvation, such as a Generalized Born approximation of the Poisson-Boltzmann equation^29^ and polarizable electrostatic terms that accommodate varying charge distributions.^30^

The first version of the Rosetta energy function was developed for proteins by Simons *et al*.^31^ Initially, it used statistical potentials describing individual residue environments and frequent residue-pair interactions derived from the Protein Databank (PDB).^32^ Later, the authors added terms for packing of van der Waals spheres, hydrogen bonding, secondary-structure, and van der Waals interactions to improve the performance of *ab initio* structure prediction.^33^ These terms were for low-resolution modeling, meaning that the scores were dependent on only the coordinates of the backbone atoms and that interactions between the side chains were treated implicitly.

To enable higher resolution modeling, in the early 2000s, Kuhlman *et al.* ^34^ implemented an all-atom energy function that emphasized atomic packing, hydrogen bonding, solvation, and protein torsion angles commonly found in folded proteins. This energy function first included a Lennard-Jones term^35^, a pairwise additive implicit solvation model,^36^ a statistically-derived electrostatics term, and a term for backbone-dependent rotamer preferences.^37^ Shortly after, several terms were added, including and an orientation-dependent hydrogen bonding term^38^ in agreement with electronic structure calculations. ^39^ This combination of traditional molecular mechanics energies and statistical torsion potentials enabled Rosetta to reach several milestones in structure prediction and design including accurate *ab initio* structure prediction.^40^ hot-spot prediction,^41, 42^ protein—protein docking,^43^ and specificity redesign^44^ as well as the first *de novo* designed protein backbone not found in nature^45^ and the first computationally designed new protein—protein interface.^46^

The Rosetta energy function has changed dramatically since it was last described in complete detail by Rohl *et al*.^47^ in 2004. It has undergone significant advances ranging from improved models of hydrogen bonding^48^ and solvation,^49^ to updated evaluation of backbone^50^ and rotamer conformations.^51^ Along the way, these developments have enabled Rosetta to address new biomolecular modeling problems including refinement of low-resolution X-ray structures,^52^ development of protein binders,^53^ and the design of vaccines,^54^ biomineralization peptides, ^55^ self-assembling materials,^56^ and enzymes that perform new functions.^57,58^ Instead of arbitrary units, the energy function is now also calibrated to compute energies in kcal/mol. The details of these energy function advances are distributed across code comments, methods development papers, application papers, and individual experts, making it challenging for Rosetta developers and users in both academia and industry to learn the underlying concepts. Moreover, members of the Rosetta community are actively working to generalize the all-atom energy function for use in different contexts^59,60^ and for all biomolecules including RNA,^61^ DNA, ^62,63^ small-molecule ligands, ^64^ non-canonical amino acids and backbones, ^65–67^ and carbohydrates, ^68^ further encouraging us to reexamine the underpinnings of the energy function. Thus, there is a need for an up-to-date description of the current energy function.

In this paper, we describe the concepts and calculations underlying the current Rosetta all-atom energy function called *beta_nov15*. Our discussion aims to expose the physical and mathematical details of the energy function required for rigorous understanding. In addition, we explain how to apply the computed energies to analyze structural models produced by Rosetta simulations. We hope this paper will provide critically needed documentation of the energy methods as well as an educational resource to help students and scientists interpret the results of these simulations.

## Computing the total Rosetta energy

The Rosetta energy function approximates the energy of a biomolecule conformation. This quantity, called ∆*E*_total_, is computed from a linear combination of energy terms *E_i_* which are calculated as a function of geometric degrees of freedom, Θ, chemical identities, aa, and scaled by a weight on each term,*w*,as shown in **Eq. 1**.

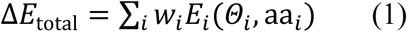

Here, we explain the Rosetta energy function term by term. First, we describe energies of interactions between non-bonded atom-pairs important for atomic packing, electrostatics, and solvation. Second, we explain empirical potentials used to model hydrogen- and disulfide-bonds. Next, we explain statistical potentials used to describe backbone and side-chain torsional preferences in proteins. After, we explain a set of terms that accommodate features not explicitly captured yet important for native structural feature recapitulation. Finally, we discuss how the energy terms are combined into a single function used to approximate the energy of biomolecules. For reference, items in the fixed width font are names of energy terms in the Rosetta code. The energy terms are summarized in**Table 1**.

**Table 1:**
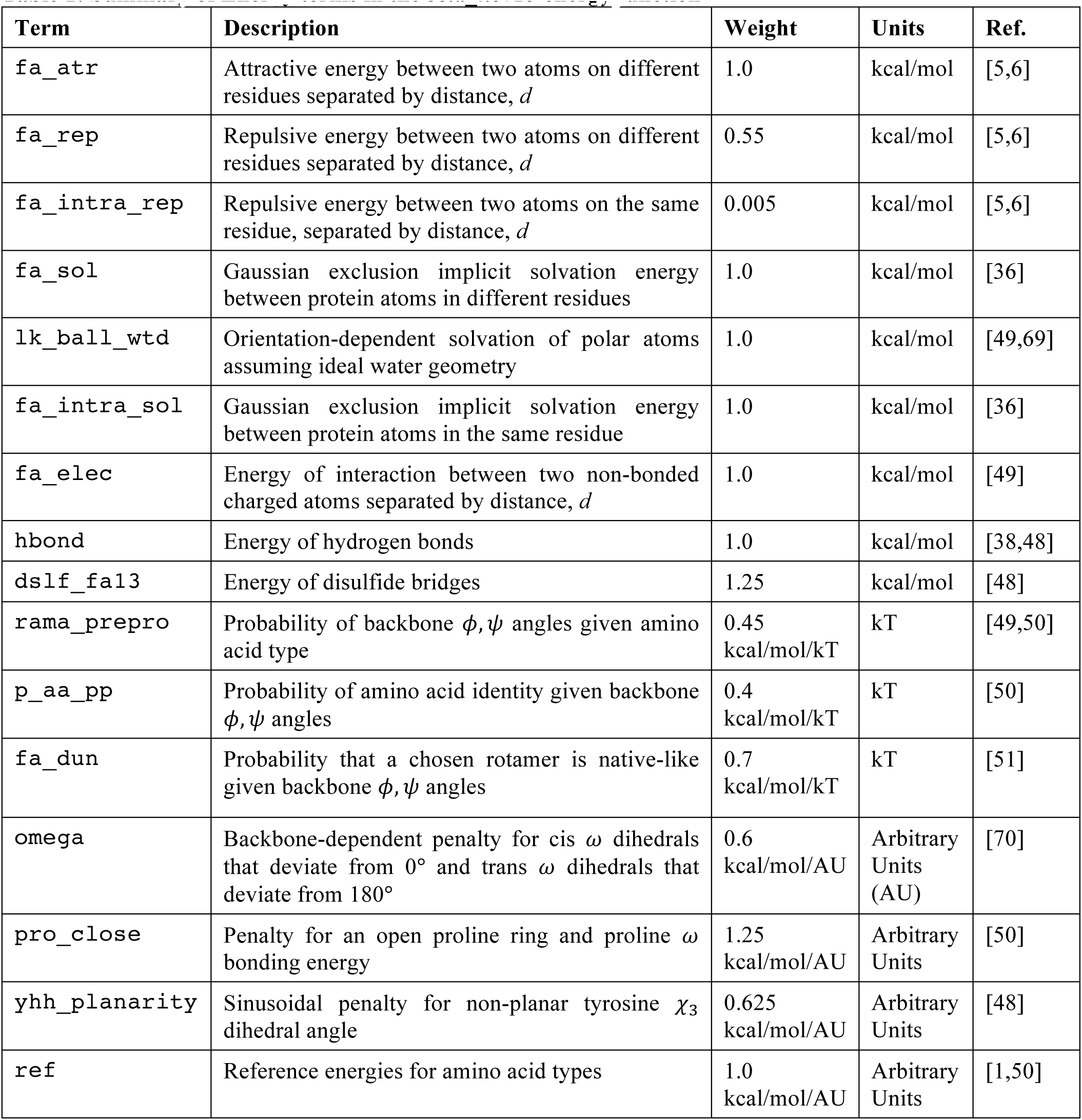
Summary of Energy terms in the *beta_nov15* energy function

*Terms for atom-pair interactions*

**van der Waals** interactions are short-range attractive and repulsive forces that vary with atom-pair distance. Whereas attractive forces result from the cross-correlated motions of electrons in neighboring non-bonded atoms, repulsive forces occur because electrons cannot occupy the same orbitals by the Pauli exclusion principle. To model van der Waals interactions, Rosetta uses the Lennard-Jones (LJ) 6-12 potential^5,6^ which calculates the interaction energy of atoms *i* and *j* in different residues given their summed atomic radii σ_*i,j*_,^a^ atom-pair distance, *d*_*i,j*_, and the geometric mean of well depths, ε*_i,j_*(**Eq. 2**). The atomic radii and well depths are derived from small molecule liquid phase data optimized in context of the energy model.^49^

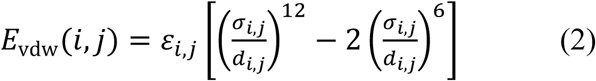

Rosetta splits the LJ potential at the function’s minimum (*d*_*i,j*_ = _σ*i,j*_) into two components that can be weighted separately: attractive (`fa_atr`) and repulsive (`fa_rep`). By decomposing the function this way, we can alter component weights without changing the minimum-energy distance or introducing any derivative discontinuities. Many conformational sampling protocols in Rosetta take advantage of this splitting by slowly increasing the weight of the repulsive component to traverse rugged energy landscapes and to prevent structures from unfolding during sampling.^71^

The repulsive van der Waals energy, `fa_rep`, varies as a function of atom-pair distance. At short distances, atomic overlap results in strong forces that lead to large changes in the energy. The steep 1/*d*_*i,j*_^12^ term can cause poor performance in minimization routines and overall structure prediction and design calculations.^72,73^ To alleviate this problem, we weaken the repulsive component by replacing the 1/*d*_*i,j*_^12^ term with a softer linear term when *d* ≤ 0.6 σ_*i,j*_. The term is computed using the atom-type specific parameters *m*_*i,j*_ and *b*_*i,j*_ which are fit to ensure derivative continuity at *d* = 0.6σ_*i,j*_ After the linear component, the function transitions smoothly to the 6-12 form until *d*_*i,j*_ =σ, where it reaches zero and remains zero (**Eq. 3; Fig. 1A**).

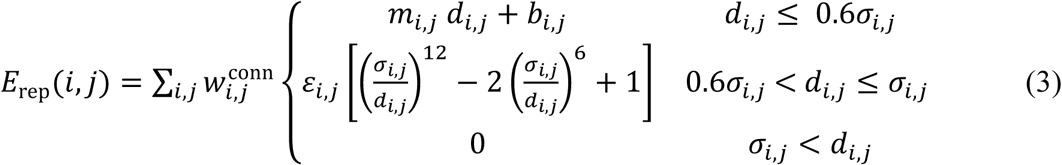

Rosetta also includes an intra-residue version of the repulsive component, `fa_intra_rep`, with the same functional form as the `fa_rep` term (**Eq. 3**)). We include this term because the knowledge-based rotamer energy (`fa_dun`, below) under-estimates intra-residue collisions.

**Figure 1:**
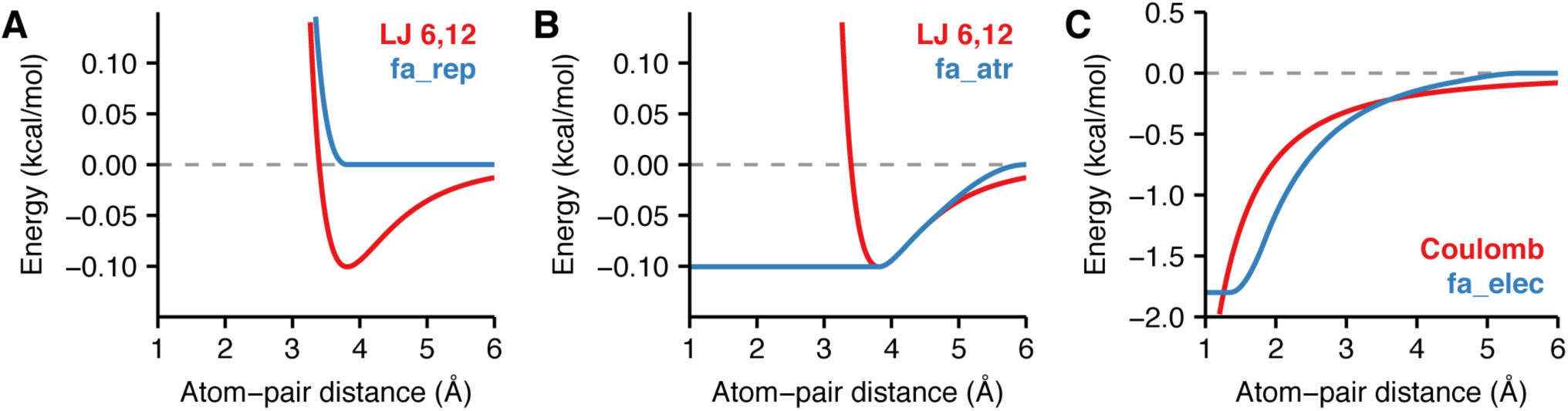
Van der Waals and electrostatics energies Comparison between pairwise energies of non-bonded atoms computed by Rosetta and the form computed by traditional molecular mechanics force fields. Here, the interaction between the backbone nitrogen and carbon are used as an example.(A) Lennard-Jones van der Waals energy with well-depths ε_Nbb_ = 0.162 and ε_Cbb_ = 0.063 and atomic radii r_Nbb_ = 1.763 and r_Cbb_ = 2.011 (red) and Rosetta `fa_rep` (blue). (B) Lennard-Jones van der Waals energy (red) and Rosetta `fa_rep` (blue). As the atom-pair distance approaches 6.0 Å, the `fa_atr` term smoothly approaches zero and deviates slightly from the original Lennard-Jones potential. (C) Coulomb electrostatics energy with a dielectric constant ε= 10, and partial charges q_Nbb_ = −0.604 and q_Cbb_ = 0.090 (red) compared with Rosetta `fa_elec` (blue). The `fa_elec` model is shifted to reach zero at the cutoff distance 6.0 Å.

The attractive van der Waals energy, `fa_atr` has a value of −ε_*i,j*_ when *d*_*i,j*_ = 0 and then transitions to the 6-12 potential as the distance increases (**Eq. 4; Fig. 1B**). For speed, we truncate the LJ term beyond 6.0 Å where the van der Waals forces are small. To avoid derivative discontinuities, we use a cubic Polynomial function, *f_poly_*(*d_i,j_*) after 4.5 Å to transition the standard Lennard-Jones functional form smoothly to zero. These smooth derivatives are necessary to ensure that bumps do not accumulate in the distributions of structural features at inflections points in the energy landscape during conformational sampling with gradient-based minimization (Sheffler 2006, Unpublished).

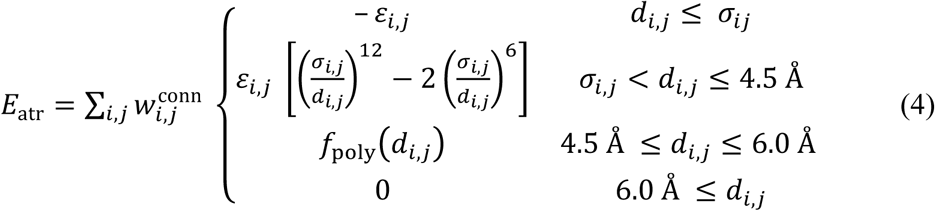

All three terms are multiplied by a connectivity weight 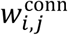 to exclude the large repulsive energetic contributions that would otherwise be calculated for atoms separated by fewer than four chemical bonds (**Eq. 5**). This weight is common to molecular force fields that assume covalent bonds are not formed or broken during a simulation. Rosetta uses four chemical bonds as the “crossover” separation when *w*_*i,j*_^conn^ transitions from zero to one (rather than the three chemical bonds used by traditional force fields) to limit the effects of double-counting due to knowledge-based torsional potentials.

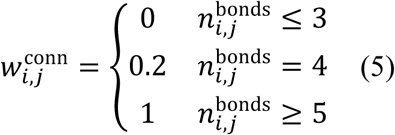

The comparison between **Eq. 2** and the modified LJ potential (**Eq. 3–4**) is shown in **Fig. 1A** and **Fig. 1B**

**Electrostatics**. Non-bonded electrostatic interactions arise from forces between fully and partially charged atoms. To evaluate these interactions, Rosetta uses Coulomb’s Law with partial charges originally taken from CHARMM and adjusted via a group optimization scheme (**Table S3**).^49^ Coulomb’s law is a pairwise term commonly expressed in terms of the distance between atoms *i* and *j* (*d*_*i,j*_), dielectric constant *ε*, partial atomic charges for each atom *q*_*i*_ and *q*_*j*_, and Coulomb’s constant, *C*_o_ = 322 Å kcal/mol *e*^-2^ (with *e* being the elementary charge) (**Eq. 6**).

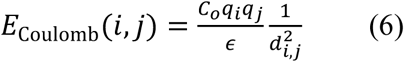

To approximate electrostatic interactions in biomolecules, we modify the potential to account for the difference in dielectric constant between the protein core and solvent-exposed surface.^74^ Specifically, we substitute the constant ε in **Eq. 6** with a sigmoidal function ε(*d*_*i,j*_ that increases from ε_core_ = 6 to ε_solvent_ = 80 when the atom-pair distance is between 0 Å and 4 Å (**Eq. 7–8**):

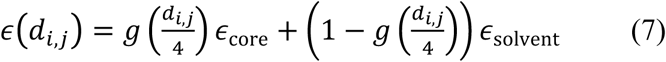

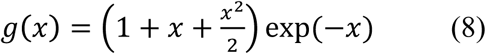

As with the van der Waals term, we make several heuristic approximations to adapt this calculation for simulations of biomolecules. To avoid strong repulsive forces at short distances, we replace the steep gradient with the constant *E*_elec_(*d*_min_) when *d*_i,j_ < 1.45 Å. Next, since the distance-dependent dielectric assumption results in dampened long-range electrostatics, for speed we truncate the potential at _max_ = 5.5 Å and we shift the Coulomb curve by subtracting a 1/d_max_^2^ term to shift the potential to zero at d_max_ (**Eq. 9**).

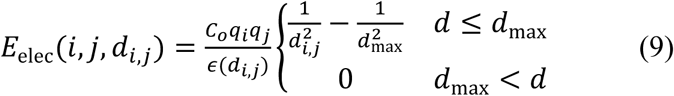

We use cubic polynomials, 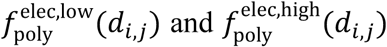 to smooth between the traditional form and our adjustments while avoiding derivative discontinuities. The energy is also multiplied by the connectivity weight, 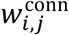(**Eq. 5**). The final modified electrostatic potential is given by **Eq. 10** and compared to the standard form in **Fig. 1C**

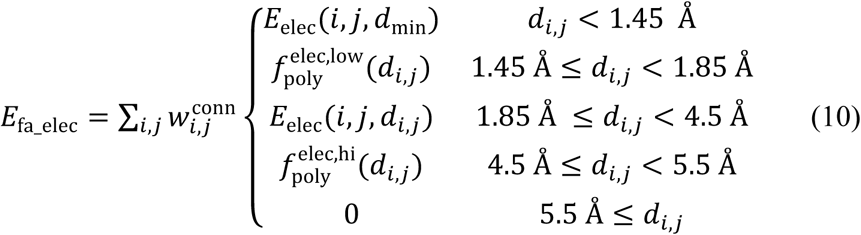

**Solvation**. Native-like protein conformations minimize the exposure of hydrophobic side chains to the surrounding polar solvent. Unfortunately, explicitly modeling all the interactions between solvent and protein atoms is computationally expensive. Instead, Rosetta represents the solvent as bulk water based upon the Lazaridis-Karplus (LK) implicit Gaussian exclusion model.^36^ Rosetta's solvation model has two components: an isotropic solvation energy, called `fa_sol`, that assumes bulk water is uniformly distributed around the atoms (**Fig. 2A**) and an anisotropic solvation energy, called lk _ball_wtd, that accounts for specific waters nearby polar atoms that form the solvation shell (**Fig. 2B**).

**Figure 2:**
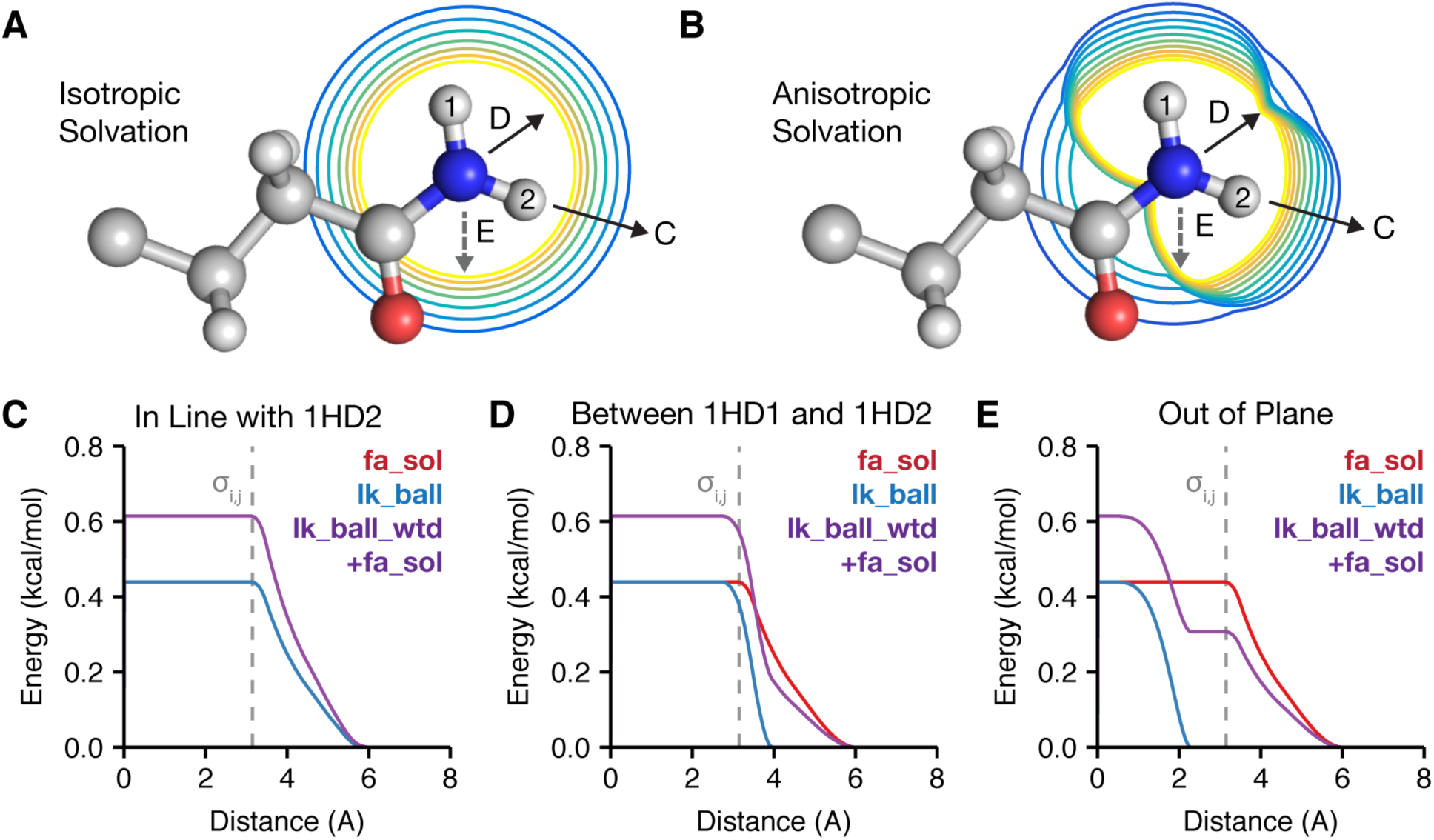
A two component Lazaridis-Karplus solvation model Rosetta uses two energy terms to evaluate the desolvation of protein side chains: an isotropic (`fa_sol`) and anisotropic (`lk_ball_wtd`) term. (A) and (B) demonstrate the difference between isotropic and anisotropic solvation of the NH2 group by CH3 on the asparagine side chain. The contours vary from low energy (blue) to high energy (yellow). The arrows represent the approach vectors for the pair potentials shown in C-E. In the bottom panel, we compare `fa_sol`, `lk_ball` and `lk _ball_wtd` energies for the solvation of the NH2 group on asparagine for three different approach angles: (C) in line with the 1HD2 atom, (D) along the bisector of the angle between 1HD1 and 1HD2 and (E) vertically down from above the plane of the hydrogens (out of plane).

The isotropic (Lazaridis-Karpus) model^36^ is based on the function *f*_desolv_ that describes the energy required to desolvate (remove contacting water) an atom *i* when approached by a neighboring atom. In Rosetta, we exclude Lazaridis-Karplus’ Δ*G*^ref^ term because we implement our own reference energy (discussed later). The energy of the atom-pair interaction varies with separation distance σ*_i,j_* experimentally determined vapor-to-water transfer free energies ∆*G*^free^, summed atomic radii 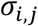 correlation length λ, and atomic volume of the desolvating atom *V*_*j*_ (**Eq. 11**).

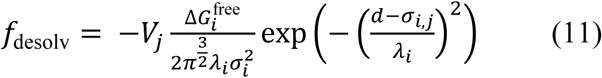

At short distances, `fa_rep` prevents atoms from overlapping; however, many protocols briefly down-weight or disable the `fa_rep` term. To avoid scenarios where *f*_desolv_ encourages atom-pair overlap in the absence of `fa_rep`, we smoothly increase the value of the function to a constant at close distances when the van der Waals spheres overlap (*d*_*i,j*_ = σ_*i,j*_). At large distances, the function asymptotically approaches zero; therefore, we truncate the function at 6.0 Å for speed. We also transition between the constants at short and long distances using distance-dependent cubic polynomials *f*_poly_^solv,low^ and *f*_poly_^solv,high^ with constants *c*_o_ = 0.3 Å and *c*_1_ = 0.2 Å that define a window for smoothing. The overall desolvation function is given by **Eq. 12**

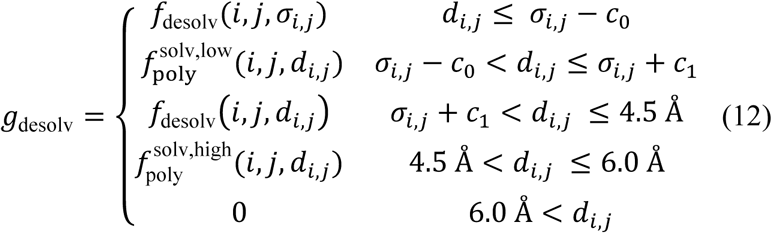

The total isotropic solvation energy (**Eq. 13**), `fa_sol`, is computed as a sum including atom *j* desolvating atom *i* and vice-versa and scaled by the previously-defined connectivity weight.

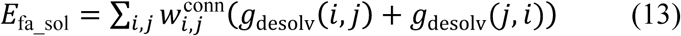

Rosetta also includes an intra-residue version of the isotropic solvation energy, `fa_intra_sol`, with the same functional form as the `fa_sol` term (Eq. 13).

A recent innovation (2016) is the addition of an energy term (`lk_ball_wtd`) to model the orientation-dependent solvation of polar atoms. This anisotropic model increases the desolvation penalty for occluding polar atoms near sites where waters may form hydrogen bonding interactions. For polar atoms, we subtract off part of the isotropic energy of **Eq. 13** and then add the anisotropic energy to account for the position of the desolvating atom relative to hypothesized water positions.

To compute the anisotropic energy, we first calculate the set of ideal water sites around atom *i*, *w*_i_ = {***v***_*i*1_,***v***_*i*2_, …}. This set contains 1 to 3 water sites, depending on the atom type of atom *i*. Each site is 2.65 Å from atom and has an optimal hydrogen-bond geometry, and we consider the potential overlap of a desolvating atom *j* with each water. The overlap is considered negligible until the van der Waals sphere of the desolvating atom *j* (radius σ_*j*_) touches the van der Waals sphere of the water at site *k* (radius σ_w_), and then the term smoothly increases over a zone of partial overlap of approximately 0.5 Å. Thus, for each water site, with coordinates ***v***_*i,k*_, we compute an occlusion measure *d*_*k*_^2^ to capture the gap between the hypothetical water and the desolvating atom *j*(**Eq. 14**), using the offset Ω = 3.7 Å^2^ to provide the ramp-up buffer.

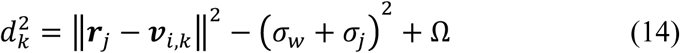

Next, we find the soft minimum of *d*_*k*_^2^ over all water sites in *w*_*i*_ by computing the log-average:

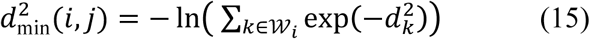

Then, *d*^2^_min_ and Ω are used to compute a damping function *f*_lkfrac_ (**Eq. 16**) that varies from zero when the desolvating atom is at least a van der Waals distance from any preferred water site to one when the desolvating atom overlaps a water site by more than ~ 0.5 Å.

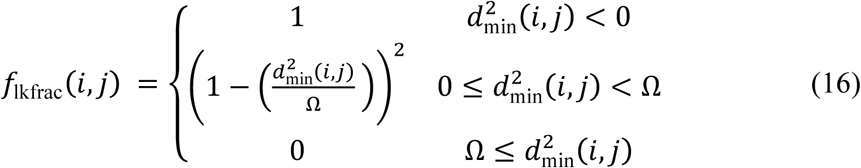

We calculate the anisotropic energy of desolvating a polar atom *E*_lk_ball_ by scaling the desolvation function *g*_desolv_ by the damping function *f*_lkfrac_ and an atom-type specific weight *w*_aniso_ that is typically ~0.7 (**Eq. 17**). The amount of isotropic solvation energy subtracted is g_desolv_ multiplied by w_iso_, where w_iso_ is an atom-type specific weight typically ~0.3 (**Eq. 18**; the total weight on the isotropic contribution through both `fa_sol` and `lk_ball_wtd` terms is thus ~0.7). The isotropic and anisotropic components are then summed to yield a new desolvation function, ℎ_desolv_ (**Eq. 19**).

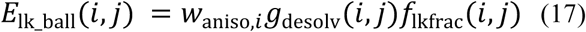

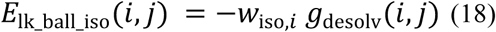

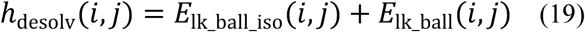

Like `fa_sol`, the energy of desolvating atom *i* by atom *j* and then *j* by *i* are summed to yield the overall `lk_ball_wtd` energy (**Eq. 20**) but only counting the desolvation of polar, hydrogen-bonding heavy atoms (O, N) defined as the set. **Fig. 2** shows a comparison between `fa_sol`, the `lk_ball` term (**Eq. 17**), and the sum of `fa_sol` and `lk_ball_wtd` for the example of an asparagine NH2 desolvated from three different approach angles. As the approach angle varies, the sum of `lk_ball` _wtd and `fa_sol` creates a larger desolvation penalty when waters sites are occluded and a smaller penalty otherwise, relative to the `fa_sol` term alone.

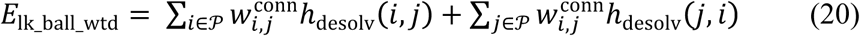

**Hydrogen bonding**. Hydrogen bonds are partially covalent interactions that form when a nucleophilic heavy atom donates electron density to a polar hydrogen.^75^ At short ranges (< 2.5 Å), they exhibit geometries that maximize orbital overlap.^76^ The interactions between hydrogen bonding groups are also partially described by electrostatics. While this hybrid covalent-electrostatic character is complex, it is crucial for capturing the structural specificity that underlies protein folding, function, and interactions.

Rosetta calculates the energy of hydrogen bonds using `fa_elec` and a term called hbond that evaluates energies based on the orientation preferences of hydrogen bonds found in high-resolution crystal structures.^38,48^ To derive this model, we curated intra-protein polar contacts from ~8,000 high resolution crystal structures (Top8000 dataset^77^) and identified features using adaptive density estimation. We then empirically fit the functional form of the energy such that the Rosetta-generated polar contacts mimic the distributions from Top8000. The resulting hydrogen bonding energy is evaluated for all pairs of donor hydrogens, *H*, and acceptors, *A*, as a function of four degrees of freedom (**Fig. 3A**): (1) the distance between the donor and acceptor, *d*_*HA*_ (2) the angle formed by the donor, acceptor, and donor-heavy atom, θ_*AHD*_ (3) the angle formed by the acceptor’s parent atom (“base”) *B*, acceptor, and the donor, θ*_BAH_* and the torsion, Φ_*B*_2_*BAH*_, formed by the donor, acceptor, and two subsequent parent atoms *B* and *B*_*2*_. (**Fig. 3A**). *B*, the parent atom of *A*, is the first atom on the shortest path to the root atom (e.g. C_α_). The *B*_*2*_ atom of *A* is the parent atom of *B* (e.g., the sp^2^ plane is defined by *B*_2_, *B*, and *A*)

**Figure 3:**
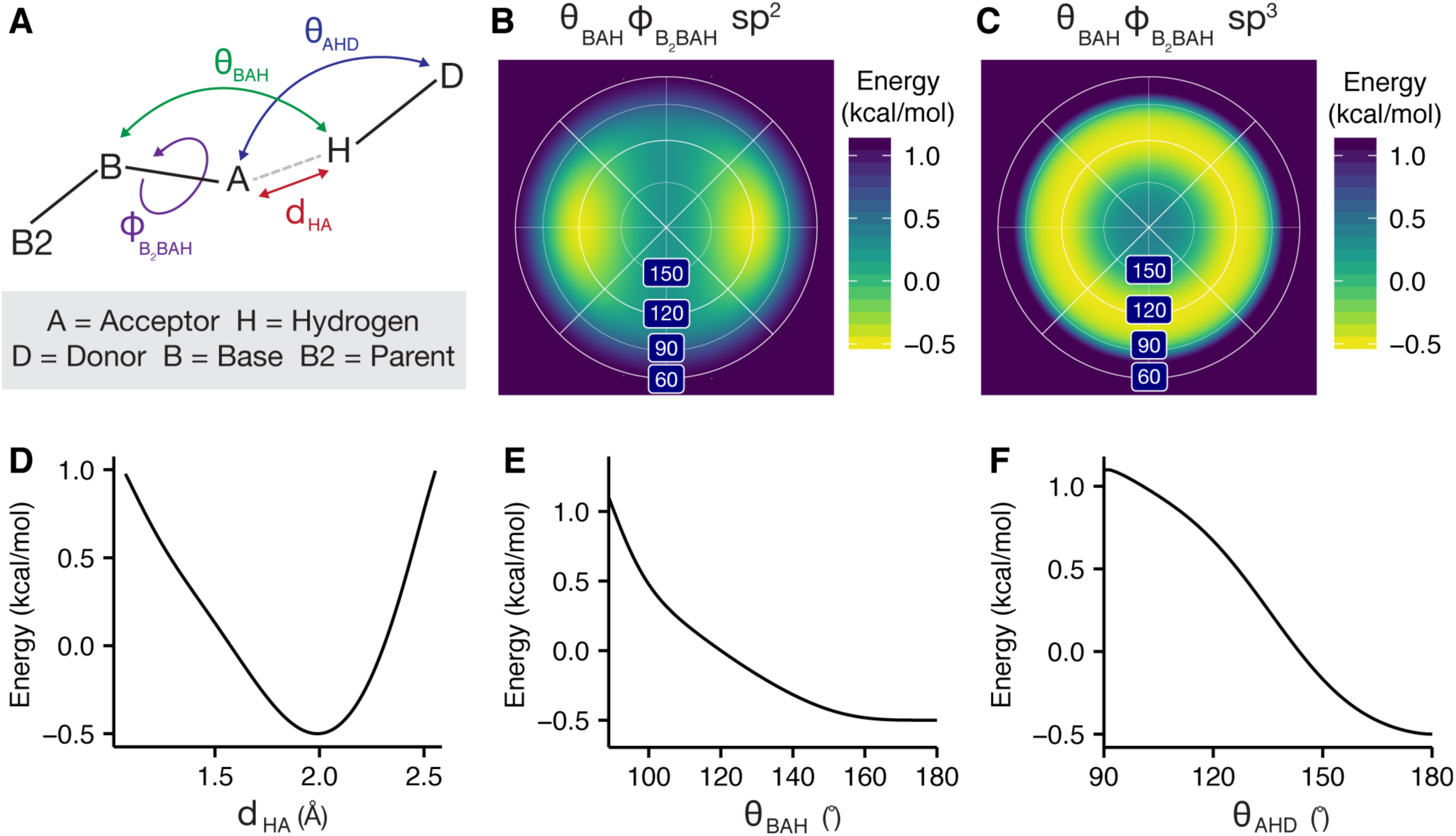
Orientation-dependent hydrogen bonding model (A) Degrees of freedom evaluated by the hydrogen bonding term: acceptor—donor distance, *d_HA_*, angle between the base, acceptor and hydrogen θ*BAH*, angle between the acceptor, hydrogen, and donor, θ_AHD_, and dihedral angle corresponding to rotation around the base—acceptor bond, ϕ_*B_2_BAH*_. (B) Lambert-azimuthal projection of the *E^B_2_BAH^*_hbond_ energy landscape for an sp^2^ hybridized acceptor.^48^ (C) *E*_hbond_*^B_2_BAH^* energy landscape for an sp^3^ hybridized acceptor. Example energies for the histidine imidazole ring acceptor hydrogen bonding with a protein backbone amide: (D) energy vs. the acceptor—donor distance, *E*^*HA*^_hbond_ (E) energy vs. the acceptor-hydrogen-donor angle, *E*^*AHD*^_hbond_ (F) energy vs. the base-acceptor—hydrogen angle, *E*^*BAH*^_hbond_.

To avoid over-counting, side-chain to backbone hydrogen bonds are excluded if the backbone group is already involved in a hydrogen bond. For speed, the component terms have simple analytic functional forms (**Fig. 3B–F**; Supporting Information **Eq. S1-7**). The term is also multiplied by two atom-type specific weights, *w*_*H*_ and *w*_*A*_, that account for the varying strength of hydrogen bonds. The overall model is given by **Eq. 21** where the *E*_hbond_^*B*_2_*BAH*^ term depends on the orbital hybridization of the acceptor,ρ. Finally, the function is also smoothed with *f*(*x*) (**Eq. 22**) to avoid derivative discontinuities and ensure that edge-case hydrogen bonds are considered.

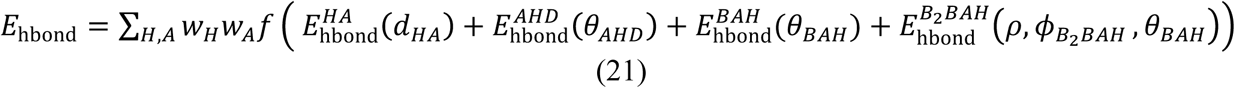

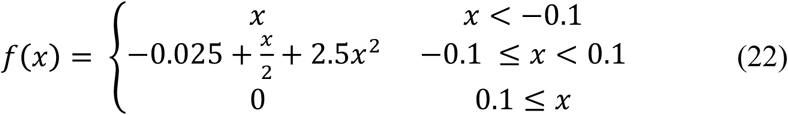

**Disulfide bonding**. Disulfide bonds are covalent interactions that link sulfur atoms in cysteine residues. Typically, in Rosetta, we rely on a tree-based kinematic system^78,79^ to keep bond lengths and angles fixed so that we may sample conformation space changing only torsions. For this reason, we do not generally need terms that evaluate bond-length and bond-angle energetics. However, with disulfide bonds and proline (below), the extra bonds cannot be represented with a tree (since a tree graph is acyclic), and thus must be treated explicitly. Thus, disulfide bonds are a special case of inter-residue covalent contact that requires a representation with more degrees of freedom. To evaluate disulfide bonding interactions, Rosetta identifies pairs of cysteines that have covalent bonds linking the Sγ atoms. Then, Rosetta computes the energy of these interactions using an orientation-dependent model called dslf _fa13.^48^ The model was derived by curating intra-protein disulfide bonds from Top8000 and identifying features using kernel density estimates. For speed, the feature distributions are modeled using skewed Gaussian functions and a mixture of 1, 2, and 3, von Mises functions (Supporting Information **Eq. S8-11**).

The overall disulfide energy is computed as a function of six degrees of freedom (**Fig. 4**) that map to four component energies. First, the geometry of the sulfur-sulfur distance *d_SS_* is evaluated by 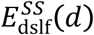. Second,the angle formed by either *C*_β1_ or *C*_β2_ with S-S bond is evaluated by 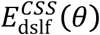 Third, the dihedral formed by either 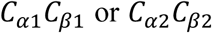 with the S-S bond is evaluated by. Finally, the dihedral formed with the S bond is evaluated by 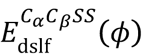. Finally, the dihedral formed by 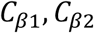 and the S-S bond is evaluated by 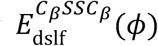. The complete disulfide bonding energy evaluated for all S-S pairs is given by **Eq. 23**

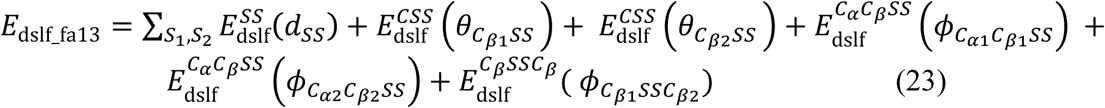

**Figure 4:**
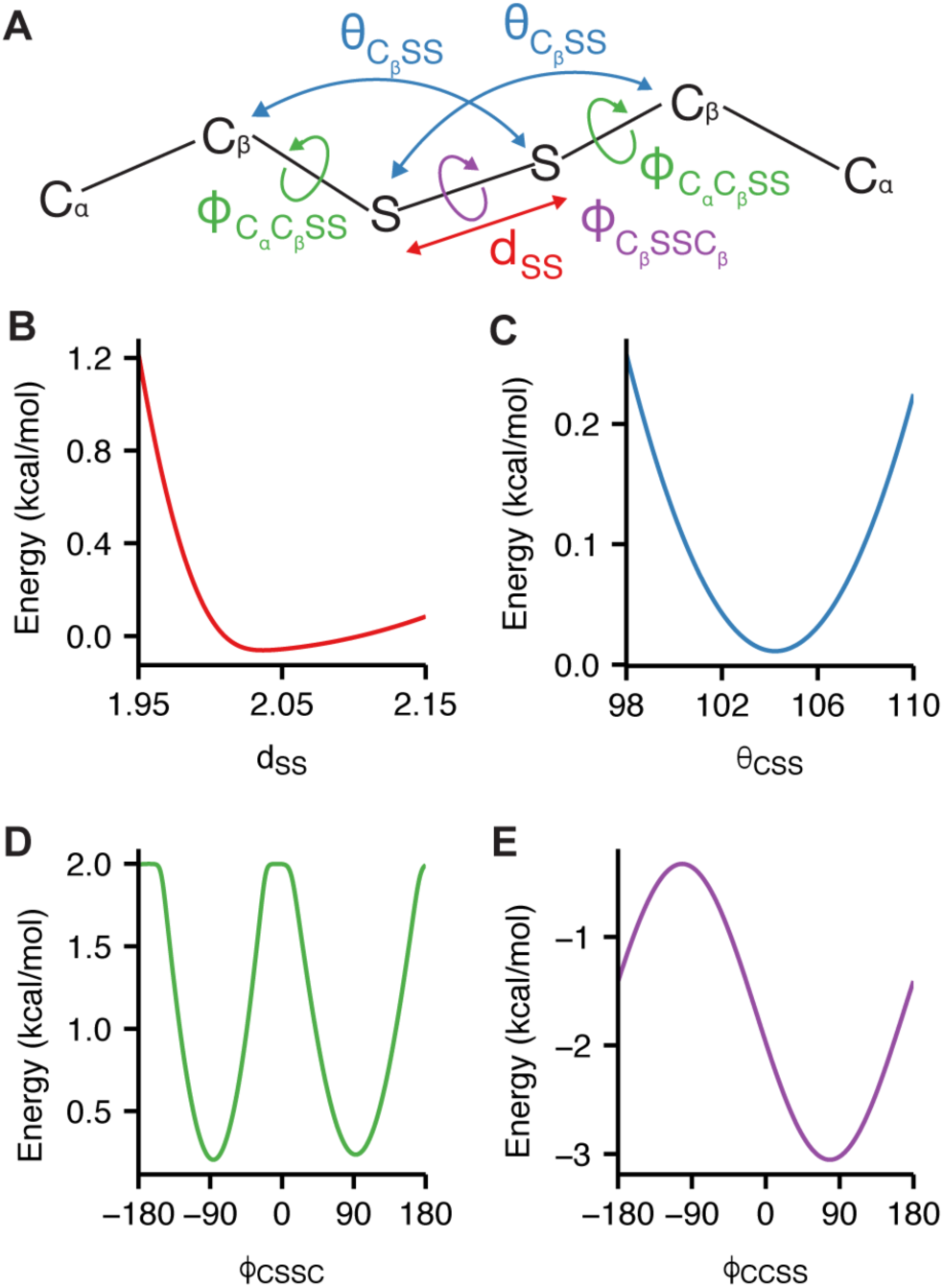
Orientation-dependent disulfide bonding model(A) Degrees of freedom evaluated by the disulfide bonding energy: sulfur—sulfur distance, angle between the β- carbon and two sulfur atoms,θ_CSS_, dihedral corresponding to rotation about the α-Carbon and sulfur bond 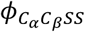 and dihedral corresponding to rotation about the 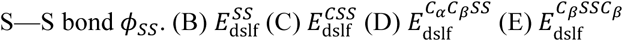

### Terms for Protein Backbone and Side Chain Torsions

Rosetta evaluates backbone and side-chain conformations in torsion space to greatly reduce the search domain and increase computational efficiency. Traditional molecular mechanics force fields describe torsional energies in terms of sines and cosines which have at times performed poorly at reproducing the observed backbone-dihedral distributions in unstructured regions.^80^ Instead, Rosetta uses several knowledge-based terms for torsion angles that are fast approximations of quantum effects and more accurately model the preferred conformations of protein backbones and side-chains.

**Ramachandran**. To evaluate backbone ϕ and ψ angles, we defined an energy term called `rama_prepro` based on Ramachandran maps for each amino acid, using torsions from 3,985 protein chains with a resolution ≤ 1.8 Å, R-factor ≤ 0.22 and sequence identity ≤ 50%.^81^ Amino acids with low electron density (in the bottom 25^th^ percentile of each residue type) were removed from the data set. The resulting ~581,000 residues were used in adaptive kernel density estimates^51^ of Ramachandran maps with a grid step of 10˚ for both ϕ and ψ. Residues preceding proline are also treated separately because they exhibit distinct ϕ,ψ preferences due to steric interactions with the proline’s C_δ_.^82^ The energy, called `rama_prepro`, is then computed by converting the probabilities to energies at the grid points via the inverted Boltzmann relation^83^ (**Eq. 24; Fig 5**). The energies are then evaluated using bicubic interpolation. The Supporting Information includes a detailed discussion of why interpolation is performed on the backbone torsional *energies* rather than the *probabilities* (**Fig. S3**, **Eqs. S12-13**).

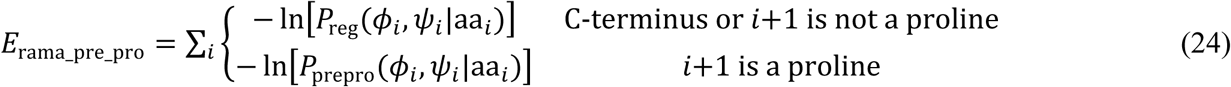

**Figure 5:**
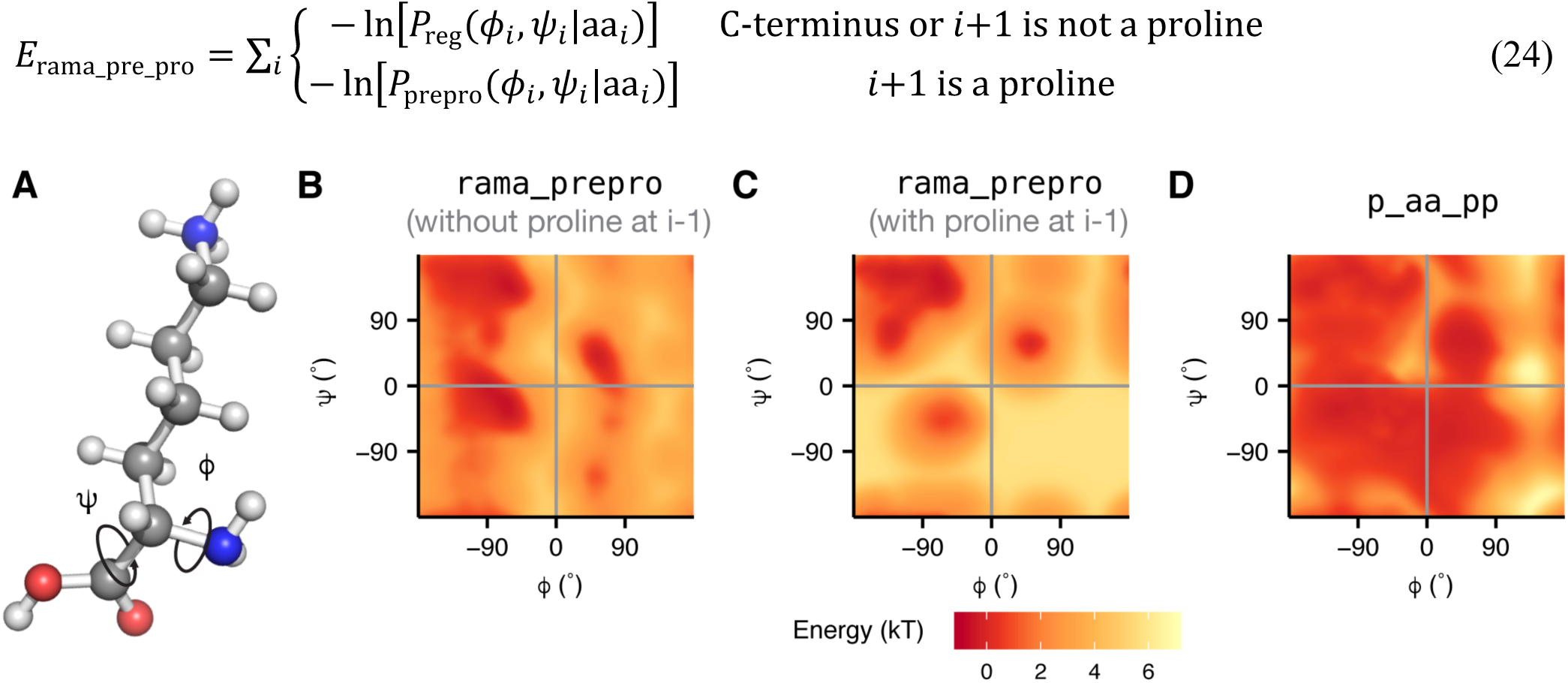
Backbone torsion energies The backbone-dependent torsion energies are demonstrated for the lysine residue. (A) The ϕ angle is defined by the backbone atoms *C*_*i*-1_−*N*−*C*_α_−*C* and the ψ angle is defined by *N*−*C*_α_−*C*−*N*_*i*+1_. (B) `rama_prepro` energy of lysine without a proline at *i*+1. (C) `rama_prepro` energy of lysine with a proline at i+1. (D) `p_aa_pp` energy of lysine.

**Backbone design term**. Rosetta also computes the likelihood of placing a specific amino acid side chain given an existing, backbone conformation. This term, called `p_aa_pp` represents the propensity of observing an amino acid relative to the other 19 canonical amino acids.^84^ The knowledge-based propensity, P(aa|ϕ,ψ), (**Eq. 25**) was derived using the adaptive kernel density estimates for (ϕ,ψ|aa) and Bayes’ rule. The equation for `p_aa_pp` is given in **Eq. 26** (**Fig. 5D**).

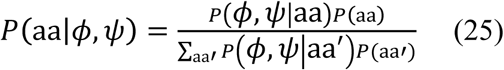

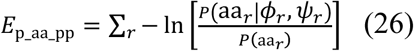

**Side-chain conformations**. Protein side chains mostly occupy discrete conformations (rotamers) separated by large energy barriers. To evaluate rotamer conformations, Rosetta derives probabilities from the 2010 backbone-dependent rotamer library (dunbrack.fccc.edu/bbdep2010), which contains the frequencies, means, and standard deviations of individual χ angles for each χ angle *k* of each rotamer of each amino acid type.^51^ The probability has three components: (1) observing a specific rotamer given the backbone dihedral angles (2) observing specific χ angles given the rotamer and (3) observing the terminal χ angle distribution, which is either Gaussian-like or continuous when the terminal χ angle is sp^2^ hybridized (**Eq. 27**). Here,*T* represents the number of rotameric χ angles +1.

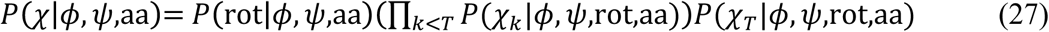

The 2010 rotamer library distinguishes between rotameric and non-rotameric torsions. A torsion is rotameric when the third of the four atoms defining the torsion is sp^3^ hybridized (i.e. preferring ~60°, ~180° and ~−60°, with steep energy barriers between the wells), If the last χ torsion is rotameric, probability p(χ*_T_*|ϕ,ψ,rot,aa) is fixed at one. On the other hand, a torsion is non-rotameric its third atom is sp^2^ hybridized: the library describes its probability distribution continuously, instead. The category of semi-rotameric amino acids with both rotameric and non-rotameric dihedrals encompasses eight amino acids: Asp, Asn, Gln, Glu, His, Phe, Tyr, and Trp.^85^

The probability of each rotamer *P*(rot|ϕ,ψ,aa) is derived from the same dataset as the Ramachandran maps described above. The probabilities were identified using adaptive kernel density estimation and the same dataset is used to estimate the mean and standard deviation for each χ dihedral in the rotamer, and µ_χ*k*_ and σ_χ*k*_, as functions of the backbone dihedrals, allowing us to compute a probability for the χ values using **Eq. 28**

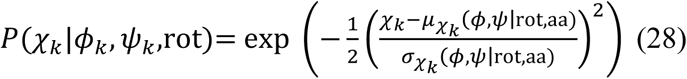

This formulation is reminiscent of the Gaussian distribution, except that it is missing the normalization coefficient of (2πσ_χ*k*_(ϕ,ψ|rot,aa))^-1/2^. After taking the log of this probability, the term resembles Hooke’s law where the spring constant is given by σ_χ*k*_^-2^(ϕ,ψ|rot,aa).

The full form of `fa_dun` is given by **Eq. 29** as a sum over all residues *r*. The difference between the rotameric and semi-rotameric models is also shown in **Fig. 6**

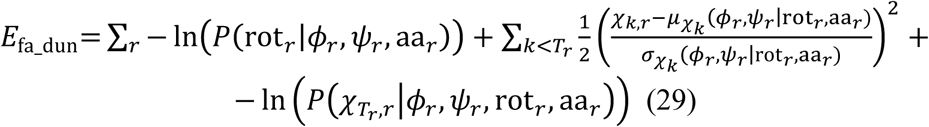

The energy from − ln 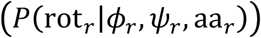 is computed using bicubic-spline interpolation; 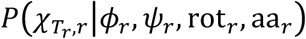 is computed using tricubic-spline interpolation. To save memory, 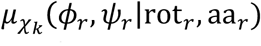 and 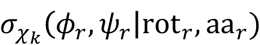 are computed using bilinear interpolation, though this has the effect of producing derivative discontinuities at the (ϕ,ψ) grid boundaries. These discontinuities, however, do not appear to produce noticeable artifacts.^50^

**Figure 6.**
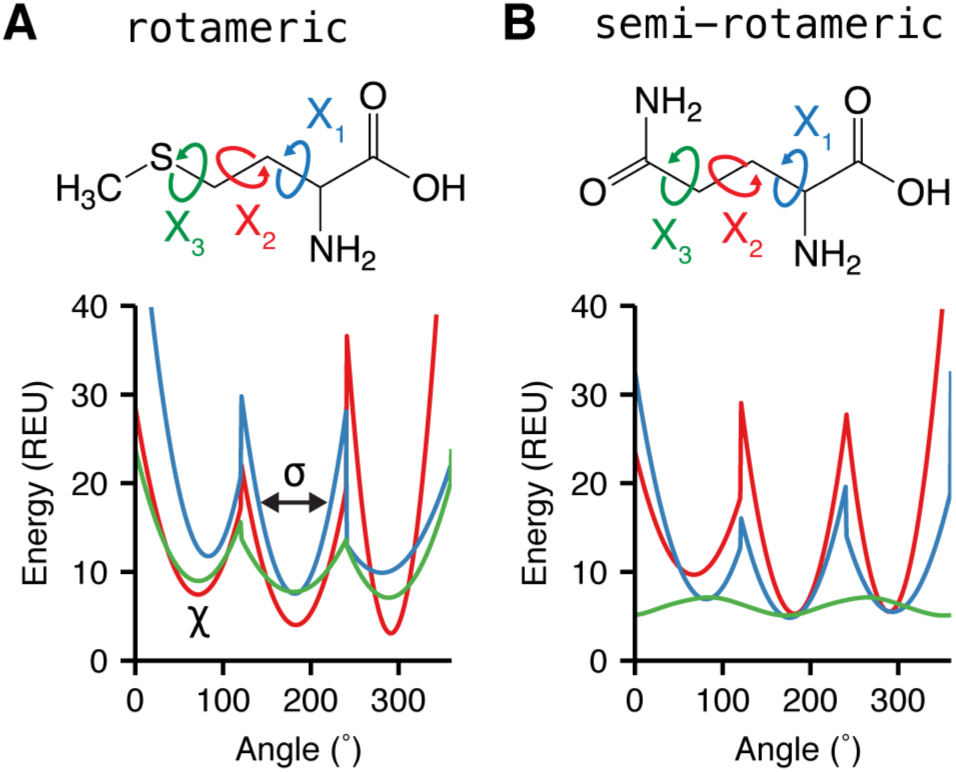
The Dunbrack rotamer energy, fa _dun, is dependent on both the ϕ and ψ backbone torsions and the χ side-chain torsions. Here, we demonstrate the variation of `fa_dun` when the backbone is fixed in an α-helical conformation with ϕ= −57˚ and ψ= −47˚, and the χ values can vary. χ_1_ is shown in blue, χ_2_ shown in red and χ_3_ shown in green. (A) χ- dependent Dunbrack energy of methionine with an *sp*^3^-hybridized terminus (B)χ-dependent energy of glutamine with an *sp*^2^-hybridized _3_ terminus. χ_1_, χ_2_ and χ_3_ of methionine and χ_1_ and χ _2_ of glutamine express rotameric behavior while χ_3_ of the latter expresses broad non-rotameric behavior.

### Terms for special case torsions

**Peptide bond dihedral angles**,ω, remain mostly fixed in a *cis-* or *trans-* conformation and depend on the backbone ϕ and ψ angles. Since the electron pair on the backbone nitrogen donates electron density to the electrophilic carbonyl carbon, the peptide bond has partial double bond character. To model this barrier to rotation, Rosetta implements a backbone-dependent harmonic penalty centered near 0° for cis and 180° for trans (**Fig. 7A**). This energy, called omega, is evaluated on all peptide bonds in the biomolecule (**Eq. 30**). The means and standard derivations of ω,µ_ω_ and σ_ω_ respectively, are backbone (ϕ,ψ) dependent, as given by kernel regressions of ω on ϕ and ψ.^70^

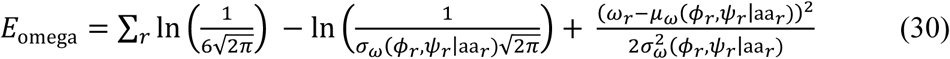

**Figure 7.**
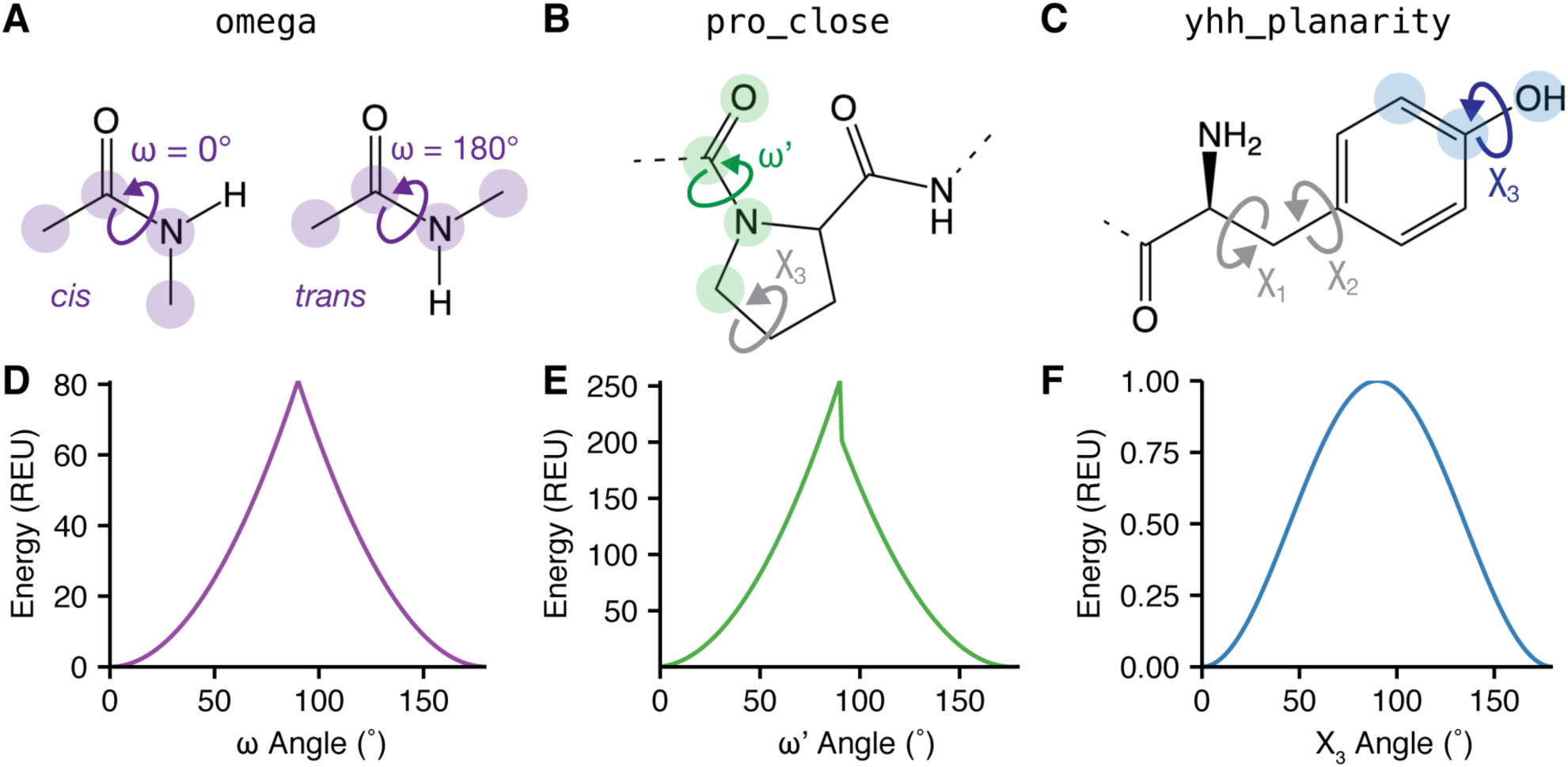
Special case torsion energies Rosetta implements three additional energy terms to model torsional degrees of freedom with acute preferences.(A) Omega torsion corresponding to rotation about C-N (B) Proline secondary omega torsion corresponding to rotation about C-N related to the C-δ in the ring. (C) Tyrosine terminal χ torsion. (D) Omega energy (E) Proline closure energy (F) Tyrosine planarity energy.

Most Rosetta protocols only search over simple torsions within chains and rigid-body degrees of freedom between chains. However, **proline’s side chain** requires special treatment because its ring cannot be represented by a kinematic tree.^86^ Therefore, Rosetta implements a proline closure term, called `pro_close` (**Fig. 7B**). There are two components to this energy, shown in**Eq. 31**. First, there is a torsional potential that operates on the dihedral formed by O_*r*- 1_–C_*r*-1_–N_*r*_–C_δ_,_*r*_, called ω_r′_ given the observed mean µ_ω′_ and standard deviation σ_ω′_, where *i* is the residue index. This term keeps the C_δ_ atom in the peptide plane. Second, to ensure correct geometry for the two hydrogens bound to C_δ_, we build a virtual atom, N_*v*_, off C_δ_ whose coordinate is controlled by χ_3_ (**Fig. 7B**). The `pro_close` term seeks to align the virtual N_*v*_ atom, directly on top of the real backbone nitrogen. The N–C_δ_–C_γ_ bond angle and the N—C_δ_ bond length are restrained to their ideal values.

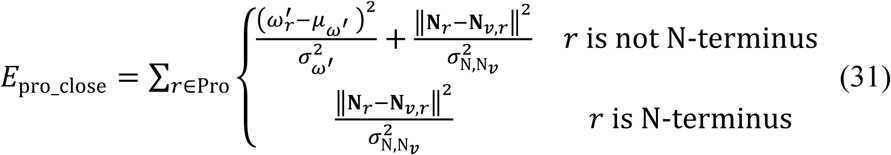

**Tyrosine** also requires special treatment for its **angle** because the hydroxyl hydrogen prefers to be in the plane of the aromatic ring.^87^ To enforce this preference, Rosetta implements a sinusoidal penalty to model the barrier to a _3_ angle that deviates from planarity. This tyrosine hydroxyl penalty is called `yhh_planarity` (**Eq. 32; Fig. 7C**).

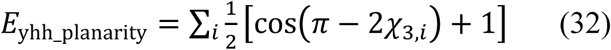

### Terms for modeling non-ideal bond lengths and angles

**Cartesian bonding energy**. Recently, modeling Cartesian degrees of freedom during gradient-based minimization has been shown to improve Rosetta’s ability to refine low-resolution structures determined by X-ray crystallography and cryo-electron microscopy,^52^ as well as its ability to discriminate near-native conformations in the absence of experimental data.^88^ These data suggest that capturing non-ideal bond lengths and angles can be important for accurate modeling of minimum-energy protein conformations. To accommodate, Rosetta now allows these “non-ideal” angles and lengths to be included as additional degrees of freedom in refinement and includes a Cartesian-minimization mode where atom coordinates are explicit degrees of freedom in optimization.

To evaluate the energetics of non-ideal bond lengths, angles and planar groups, an energy term called `cart_bonded` represents the deviation of these degrees of freedom from ideal using harmonic potentials (**Eq. 32–34**). Here, *d_i_* is a bonded-atom-pair distance with *d_i,0_*, as its ideal distance, θ*_i_* is a bond angle with θ_*i,0*_ as its ideal angle, and ϕ*_i_* is a bond torsion or improper torsion with ϕ*_i,0_*, as its ideal value and ρ*_i_* as its periodicity. The ideal bond lengths and angles^89,90^ were selected based on their ability to rebuild side chains observed in crystal structures (Kevin Karplus & James J. Havranek, unpublished); they were subsequently modified empirically.^50^ The spring constants for the angle and length terms are from CHARMM32.^19^ Finally, all planar groups and the C_β_ “pseudo-torsion” are constrained using empirically derived values and spring constants:

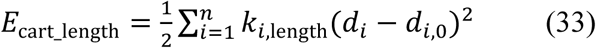

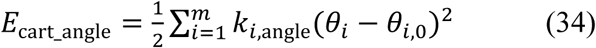

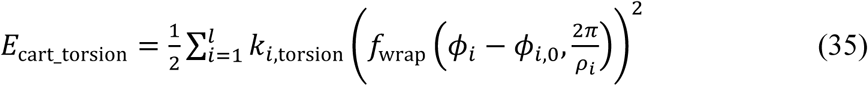

The function *f*_wrap_(*x, y*) wraps x to the range [0,y). To avoid double counting in the case of *E*_carttorsion_ the spring constant k_*i*,torsion_ is zero when the torsion ϕ*_i_* is being scored by either the rama or fadun terms

### Terms for Protein Design

**Design reference energy**. The terms above are sufficient for comparing different protein conformations with a fixed sequence. However, protein design simulations compare the relative stability of different amino acid sequences given a desired structure to identify models that exhibit a large free energy gap between the folded and unfolded states. Explicit calculations of unfolded state free energies are computationally expensive and error prone. Rosetta therefore approximates the relative energies of the unfolded state ensembles using an unfolded state reference energy, called ref.

Rosetta calculates the reference energy as a sum of individual constant unfolded state reference energies, ∆ *G*_*i*_^ref^, for each amino acid, aa_*i*_(**Eq. 36**).^1^

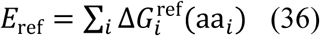

The ∆*G*_*i*_^ref^ values are empirically optimized by searching for values that maximize native sequence recovery (discussed below) during design simulations on a large set of high-resolution crystal structures.^49,50^ During design, this energy term helps normalize the observed frequencies of the different amino acids. When design is turned off, the term contributes a constant offset for a fixed sequence.

### Bringing the energy terms together

The Rosetta energy function combines all the terms using a weighted linear sum to approximate free energies (**Table 1**). Historically, we adjust the weights and parameters to balance the energetic contribution from each term. This balance is important because the van der Waals, solvation, and electrostatics energies partially capture torsional preferences and overlap can cause errors as a result of double counting atomic or residue specific contributions.^91^ More recently, we fix physics-based terms with weights of 1.0 and perturb other weights and atomic-level parameters using a Nelder-Mead^92^ scheme to optimize agreement of Rosetta calculations with small-molecule thermodynamic data and high resolutions structural features.^49^ The energy function parameters have evolved over the years by optimizing the performance of multiple scientific benchmarks (**Table 2**).^49,50,93^ These benchmarks were chosen to test recovery of native-like structural features, ranging from individual hydrogen bond geometries to thermodynamic properties and interface conformations. In addition, and more recently, Song *et al*.,^94^ Conway *et al*.^95^ and O’Meara *et al*.^46^ have fit intra-term parameters to recover features of the experimentally determined folded conformations. An in-depth review of energy function benchmarking can be found in Leaver-Fay *et al.*^96^**Table S3** lists the Rosetta database files containing the current full set of physical parameters for each score term.

**Table 2:** Common energy function benchmarking methods

| Test | Description | Ref. |
| --- | --- | --- |
| Sequence Recovery | Percentage of the native sequence recovered after backbone redesign | [1,50] |
| Rotamer Recovery | Percentage of native rotamers recovered after full repacking | [50] |
| $\Delta\Delta G$ Prediction | Prediction of free energy changes upon mutation | [97] |
| Loop Modeling | Prediction of loop conformations | [98] |
| High-resolution refinement | Discrimination of native-like decoys upon refinement of *ab initio* protein models | [99] |
| Docking | Prediction of protein-protein interfaces | [100,101] |
| Homology Modeling | Structure prediction incorporating homologous information from templates | [102] |
| Thermodynamic properties | Recapitulation of thermodynamic properties of protein side-chain analogues | [17] |
| Recapitulation of Xtal structure geometries | Recapitulation of features (e.g. atom-pair distance distribution) from high-resolution protein crystal structures | [49] |

### Energy Function Units

Initially, Rosetta energies were expressed in a generic unit, called the Rosetta Energy Unit (REU). This choice was made because some original terms of Rosetta energy were not in kcal/mol, and the use of statistical potentials convoluted interpretation of the energy. Over time, the physical meaning of Rosetta energies has been extensively debated within and outside the community, and several steps have been taken to clarify interpretation. The current energy function (*`beta_nov`15*) was parameterized on small molecule thermodynamic data and high-resolution protein structures in units of kcal/mol.^49^ The optimization data show a strong correlation (*R* = 0.994) between the experimental data and values predicted by Rosetta ( ΔΔ*G* upon mutation, small molecule Δ*H*_vap_; **Fig. S2**); therefore, as is standard practice for molecular force fields such as OPLS, CHARMM, and AMBER, we now express energies in kcal/mol.

## Energies in action: Using individual energy terms to analyze Rosetta models

Rosetta energy terms are mathematical models of the physics that governs protein structure, stability, and association. Therefore, the decomposed relative energies of a structure or ensemble of structures can expose important details about the biomolecular model. Now that we have presented the details of each energy term, we here demonstrate how energies can be applied to detailed interpretations of structural models. In this section, we discuss two common structure calculations: (1) estimating the free energy change (∆∆*G*) of mutation^97^ and (2) modeling the structure of a protein-protein interface.^101^

**ΔΔ*G* of mutation**. The first example demonstrates how Rosetta can be used to estimate and rationalize thermodynamic parameters. Here, we present an example ∆∆*G* of mutation calculation for the T193V mutation in the RT-RH derived peptide bound to HIV-1 protease (PDB 1kjg, **Fig. 8A**).^103^ The details of this calculation are provided in the Supporting Information.

**Figure 8:**
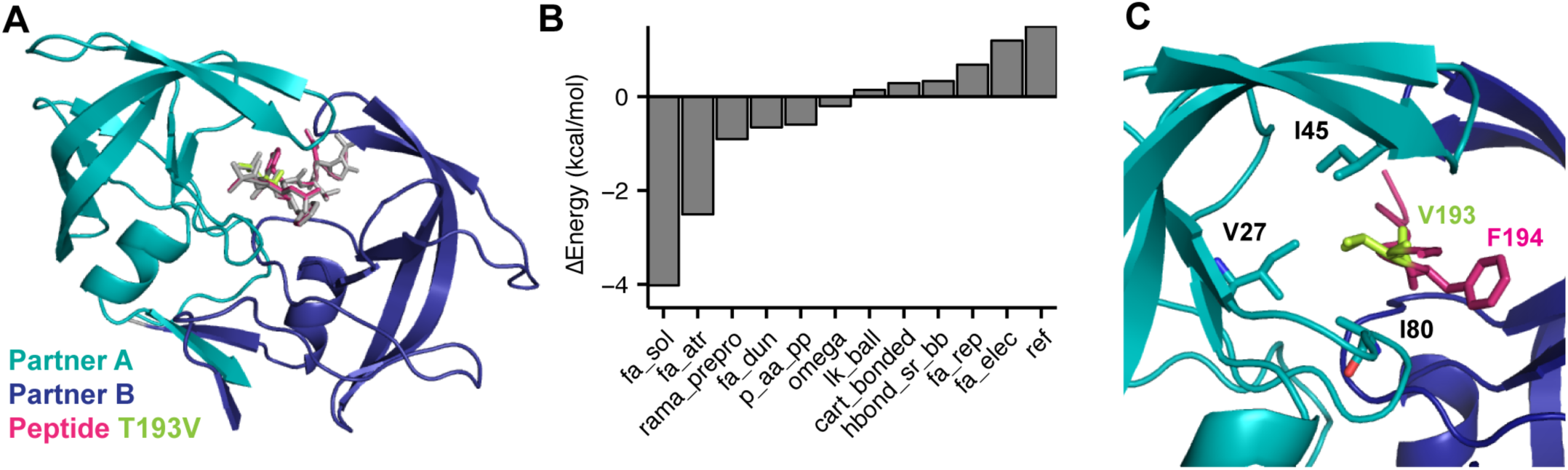
Structural model of the HIV-1 protease bound to the T4V mutant RT-RH derived peptide Structural model of the native HIV-1 peptidase (teal and dark blue), bound to the native peptide (gray) superimposed onto the T4V mutant peptide (magenta). (B) Contributions greater than + 0.1 kcal/mol to the ∆∆*G* of mutation for T4V. The remaining contributions are: `dslf_fa`13 = 0 kcal/mol, `hbond_lr_bb` = −0.09 kcal/mol, `hbond_bb_sc` = −0.05, `hbond_sc` = −0.0104, `fa_intra_rep` = 0.01, `fa_intra_sol` = −0.07, and `yhh_planarity` = 0. (C) Hydrophobic patch of residues surrounding position four on the RT-RH peptide.

Rosetta calculates the ∆∆*G* of the T193V mutation to be −4.95 kcal/mol, and the experiment^103^ measured −1.11 kcal/mol. Both the experiment and calculation reveal that T193V is stabilizing: yet, these numbers alone do not reveal which specific interactions are responsible for the stabilization. To investigate, we used various analysis tools accessible in PyRosetta^104^ to identify important energetic contributions to the total ∆∆*G*. First, we decomposed the ∆∆*G* into individual energy terms and observe the balance of terms, both favorable and unfavorable, that sum to the total (**Fig. 8B**). To decompose the most favorable term, ∆`fa_sol`, we used the `print_residue`_ pair _energies function to identify residues that interact with the mutation site (in this case, residue 4) to produce a nonzero residue pair solvation energy. With the resulting table, we found a hydrophobic pocket around the mutation site formed by residues V27, I45, G46, and I80 on HIV peptidase and residue F194 on the peptide made a large (> 0.05 kcal/mol) and favorable contribution to the change in solvation energy (**Fig. 8C**).

We further investigated this result on the atomic level with the function `print_atom`_`pair_energy_table` by generating atom-pair energy tables (Supporting Information) for residues 5, 27, 45, 46, and 80 against both threonine and valine at residue 193 (Example for residue 80 in **Table 3**). Here, we find that the specific substitution of the polar hydroxyl on threonine with nonpolar alkyl group on valine stabilizes the peptide in the hydrophobic protease pocket. This result is consistent with chemical intuition and demonstrates how breaking down the total energies can provide insight into characteristics of the mutated structures.

**Table 3:** Change in atom pair energies between I80 and T4 versus V4 in kcal/mol

| T193→V193<br>Atoms | I80 Atoms |  |  |  |
| --- | --- | --- | --- | --- |
|  | CB | CG1 | CG2 | CD1 |
| N | 0.000 | 0.000 | 0.000 | 0.000 |
| CA | 0.000 | 0.000 | 0.000 | 0.004 |
| C | 0.000 | 0.000 | 0.000 | 0.008 |
| O | 0.000 | 0.000 | 0.000 | -0.010 |
| CB | 0.000 | 0.054 | 0.000 | -0.002 |
| OG1 → CG1 | 0.008 | -0.054 | -0.316 | -0.398 |
| CG2 → CG2' | 0.000 | 0.000 | 0.001 | 0.020 |

**Protein-protein docking**. The second example shows how the Rosetta energies of an ensemble of models can be used to discriminate between models and investigate the characteristics of a protein–protein interface. Below, we investigate docked models of West Nile Virus envelope protein and a neutralizing antibody (PDB 1ztx; **Fig. 9A**).^105^ Calculation details can be found in the Supporting Information.

**Figure 9:**
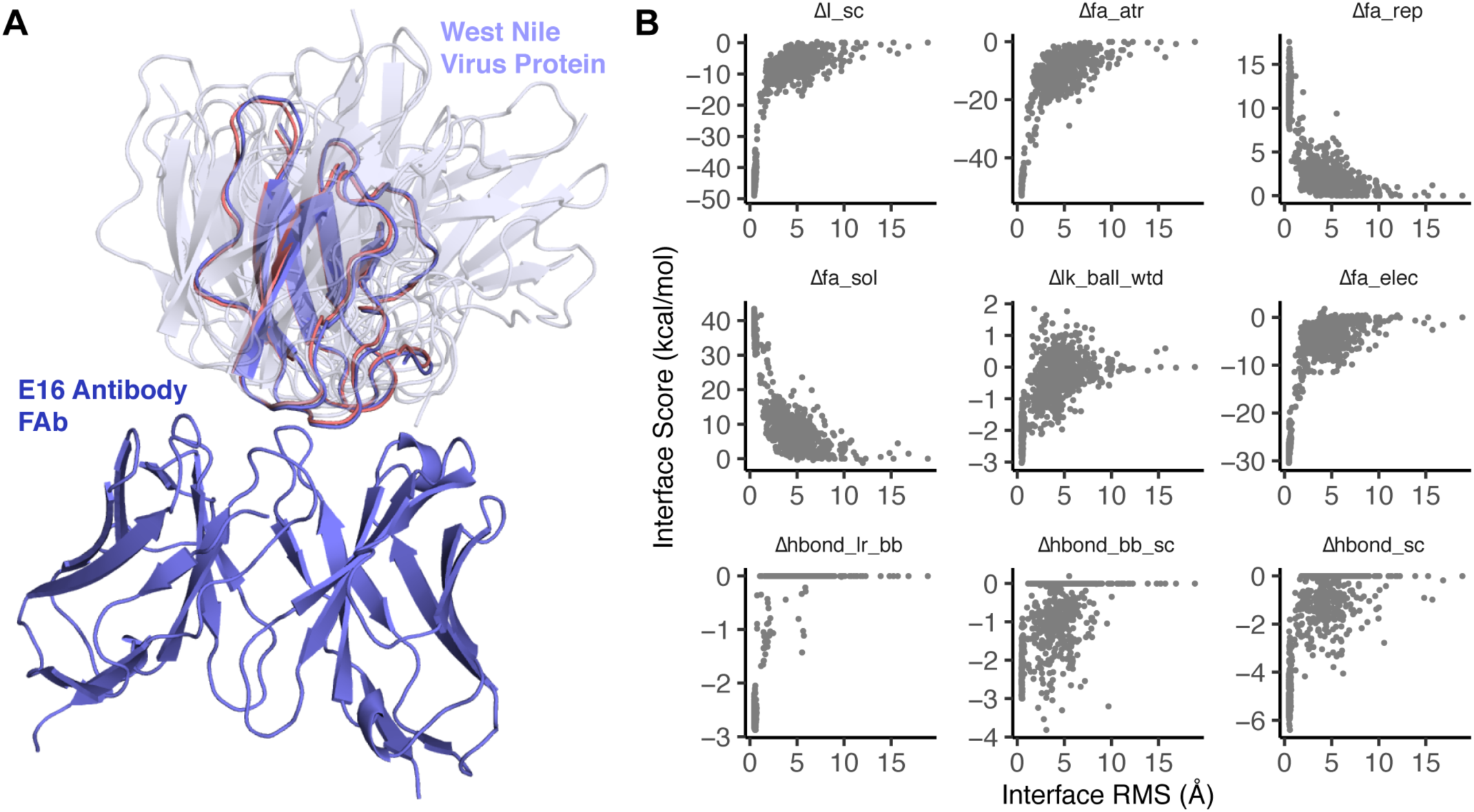
Using energies to discriminate docked models of West Nile Virus and the E16 neutralizing antibody (A) Comparison of the native E16 antibody (purple) docked to the lowest RMS model of the West Nile Virus envelope protein and several other random models of varying energy to show sampling diversity (gray, semi transparent). (B) Change in the interface energy relative to the unbound state versus RMS to native. Models at low RMS to the native interface have a low overall interface energy due to favorable van der Waals contacts, electrostatic interactions, and side-chain hydrogen bonds, as reflected by the Δ`fa_atr`,Δ`fa_elec`, and Δ`hbond_sc` energy terms.

To evaluate the docked models, we examine the variation of energies as a function of the root mean squared deviation (RMS) between the residues at the interface in each model and the known structure. For our calculation, interface residues are residues with a C_β_ atom less than 8.0 Å away from the C_β_ of aresidue in the other docking partner. The plot of energies against RMS values is called a *funnel plot* and is intended to mimic the funnel-like energy landscape of protein folding and binding.

Like the previous example, we decompose the energies to yield information about the nature of interactions at the interface. Here, we observed significant changes in the following energy terms upon interface formation relative to the unbound state: `fa_atr`, `fa_rep`, `fa_sol`, `lk_ball_wtd`, `fa_elec`, `hbond_lr_bb`, `hbond_bb_sc`, and hbond _sc (**Fig. 9B**). Change in the Lennard-Jones energy upon interface formation is due to the introduction of atom-atom contacts at the interface. As more atoms come into contact near the native conformation (RMS→0), the favorable, attractive energy (fa _atr) decreases whereas the unfavorable, repulsive energy (Δ`fa_rep`) increases. Change in the isotropic solvation energy (`fa_sol`) is positive (unfavorable), indicating that upon interface formation, polar residues are buried. Balancing the desolvation penalty, the change in polar solvation energy (`lk_ball` _wtd) and electrostatics (`fa_elec`) is negative due to polar contacts forming at the interface. Finally, the three hydrogen bonding energies (`hbond_lr`_ bb, `hbond_bb_sc`, and `hbond_sc`) reflect the formation of backbone—backbone, backbone–side-chain, and side-chain–side-chain hydrogen bonds at the interface.

## Discussion

The Rosetta energy function represents our collaboration’s ongoing pursuit to model the rules in nature that govern biomolecular structure, stability, and association. This paper summarizes the latest version which brings together fundamental physical theories, statistical mechanical models, and observations of protein structures. This work represents almost 20 years of interdisciplinary collaboration in the Rosetta community, which in turn builds on and incorporates decades of work outside the community.

After 20 years, we have improved physical theories, structural data, representations, experiments, and computational tools; yet, energy functions are far from perfect. Compared to the first torsional potentials, energy functions are also now vastly more complex. There are countless ways to arrive at more accurate energy functions. Here, we discuss grand challenges specific to development of the Rosetta energy function in the coming decade.

### Modeling biomolecules other than proteins

The Rosetta energy function was originally developed to predict and design protein structures. A clear artifact of this goal is the energy function’s dependence on statistical potentials derived from protein X-ray crystal structures. Today, the Rosetta community also pursues goals ranging from design of synthetic macromolecules to predicting interactions and structures of other biomolecules such as glycoproteins and RNA. Accordingly, an active research thrust is to generalize the all-atom energy function for all biomolecules.

Many of the physically-derived terms (e.g. van der Waals) have already been made compatible with non-canonical amino acids and non-protein biomolecules (**Table S5**). Recently, Bhardwaj, Mulligan & Bahl *et al.*^67^ adapted the `rama_prepro`, `p_aa_pp`, `fa_dun`, `pro_close`, omega, `dslf_fa`13, `yhh_planarity` and ref terms to be compatible with mixed-chirality peptides. Several of Rosetta’s statistical potentials are validated against quantum mechanical calculations for evaluating for non-protein models (**Table 4**). The first non-protein terms were added by Havranek *et al.*^106^ and Yu *et al.*^107^ who modified the hydrogen bonding potential to capture planar hydrogen bonds between protein side chains and nucleic acid bases. Renfrew *et al.*^65,108^ added molecular mechanics torsions and Lennard-Jones terms to model proteins with non-canonical amino acids, oligosaccharides,peptides, and oligo-peptoids.^66^ Labonte *et al*^68^ implemented Woods’ CarboHydrate-Intrinsic (CHI) function^109,110^ which evaluates glycan geometries given the axial-equatorial character of the bonds. In addition, Das *et al.* added a set of terms to model Watson-Crick base pairing, π-π interactions in base stacking, and torsional potentials important for predicting and designing RNA structures.^61,111–113^ These terms are presented in detail in the Supporting Information.

**Table 4:**
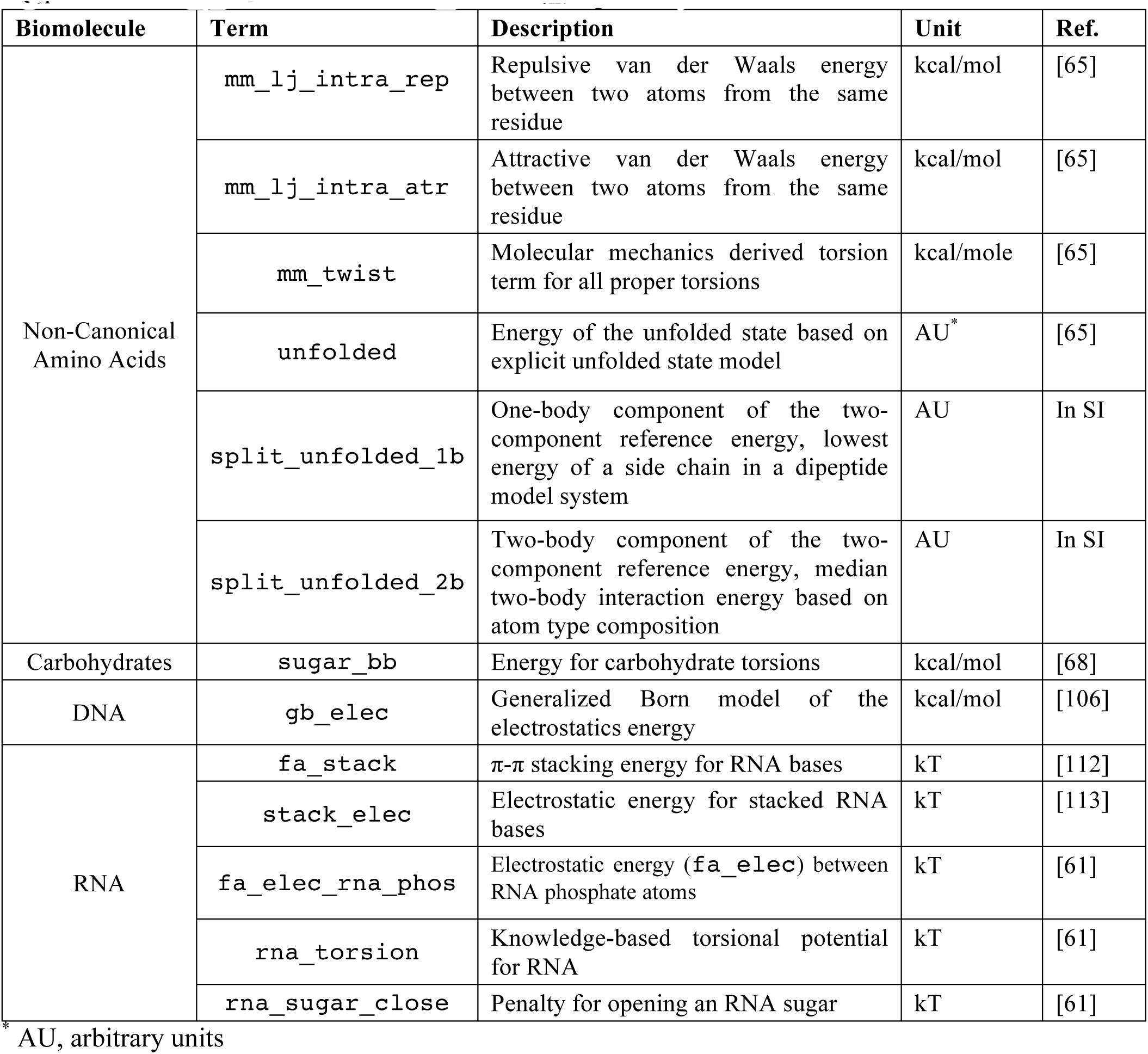
New energy terms for biomolecules other than proteins

Expanding Rosetta’s chemical library brings new challenges. Currently, there are separate energy functions for various types of biomolecules. Typically, these functions mix physically-derived terms from the protein energy function with molecule-specific statistical potentials, custom weights, and possibly custom atomic parameters. If nature only uses one energy function, why do we need so many? Some discrepancies may result from features that we do not model explicitly, such as π-π, n-π^∗^ and cation-interactions. Efforts to converge on a single energy function will therefore pose interesting questions about the set of universal physical determinants of biomolecular structure.

AU, arbitrary units

### Capturing the intra and extra-cellular environment

Rosetta traditionally models the solvent surrounding the protein using the Lazaridis-Karplus (LK) model, which assumes a solvent environment made of pure water. In contrast, biology operates under various conditions influenced by pH, redox potential, temperature, solvent viscosity, chaotropes, kosmotropes, and polarizability. Therefore, modeling more details of the intra and extra-cellular environment would enable Rosetta to identify structures important in different biological contexts.

Currently, Rosetta includes two groups of energy terms to model alternate environments (**Table 5**). Kilambi *et al.*^59,114^ implemented a method to account for alternate protonation states due to pH changes. In addition, Rosetta implements Lazaridis’ Implicit Membrane Model for modeling proteins in a lipid bilayer environment.^36,60,115,116^ While these improve structure prediction accuracy, both models require more computation time. This trade-off between the need for detail and computational complexity will be evaluated as Rosetta aims to model more complicated biological systems and contexts.

**Table 5:**
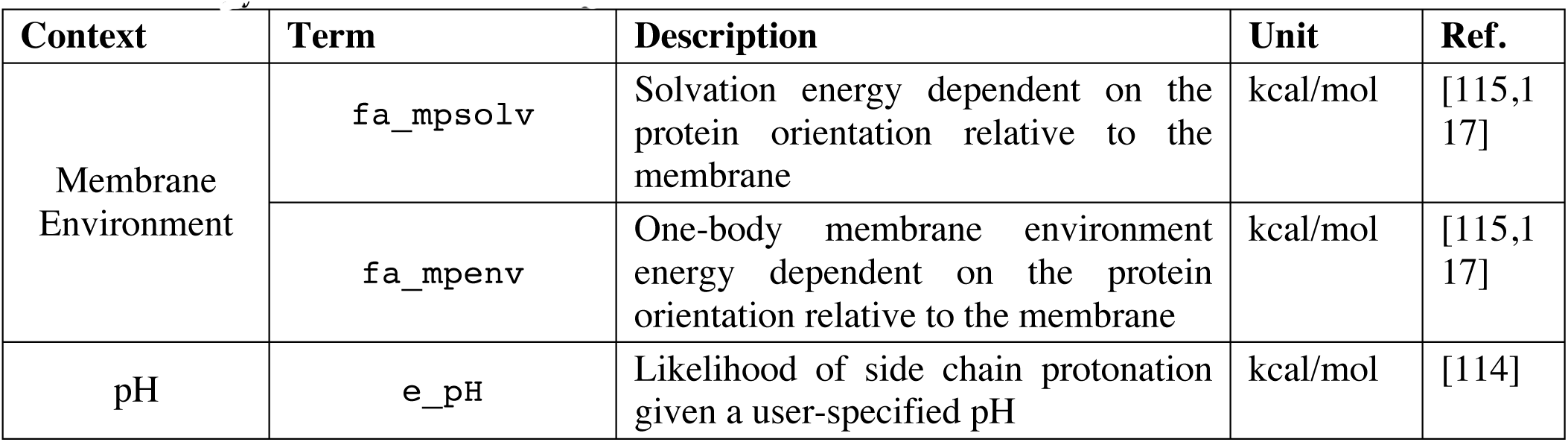
Energy terms for structure prediction in different contexts

### The origin of energy models: top-down versus bottom-up development

Traditionally, energy functions are developed using a bottom-up approach: experimental observables serve as building blocks to parameterize physics-based formulas. The advent of powerful optimization techniques and artificial intelligence recently empowered the top-down category where numerical methods are used to derive models and/or parameters. Top-down approaches have been used to solve problems in various fields including structural biology and bioinformatics. Recently, top-down development was also applied to optimizing the Lennard-Jones, Lazaridis-Karplus, and Coulomb parameters in the Rosetta energy function (parameters in **Table S4-S6**).^49,92^

Top-down approaches have enormous potential to improve the accuracy of biomolecular modeling because more parameters can vary and the objective function can be minimized with more benchmarks. These approaches also introduce new challenges. With any computer-derived models, there is a risk of over-fitting as validation via structure prediction datasets reflect observable states, whereas simulations are intended to predict features of states that experiments cannot yet observe. Computer-derived parameters also introduce a unique kind of uncertainty. Consider the following scenario: the performance of scientific benchmarks improves as physical atomic parameters are perturbed away from the measured experimental values. As there is less physical-basis for parameters, are the predictions and interpretations still meaningful?

Top-down development will also provide power to develop more complicated energy functions. Currently, the Rosetta energy function advances by incrementally addressing weaknesses: with each new paper, we modify analytic formulas, add corrective terms, and adjust weights. As this paper demonstrates, the energy function is significantly more complicated than the initial theoretical forms. Given this complexity increase, an interesting approach to leverage the power of top-down development would be to simplify and subtract terms to evaluate individual benefits.

### highly interdisciplinary endeavor

The Rosetta energy function has advanced rapidly due to the Rosetta Community: a highly-interdisciplinary collaboration between scientists with diverse backgrounds located in over 50 labs around the world. The many facets of our team enable us to probe different aspects of the energy function. For example, expert computer scientists and applied mathematicians have implemented algorithms to speed up calculations. Dedicated software engineers maintain the code and maintain a platform for scientific benchmark testing. Physicists and chemists develop new energy terms that better model the physical rules found in nature. Structural biologists maintain a focus on created biological features and functions. We look forward to leveraging this powerful interdisciplinary scientific team as we head into the next decade of energy function advances.

## Conclusion: A living energy function

For the first time since 2004,^47^ we have documented all of the mathematical and physical details of the Rosetta all-atom energy function highlighting the latest upgrades to both the underlying science and the speed of calculations. In addition, we illustrated how the energies can be used to analyze output models from Rosetta simulations. These advances have enabled Rosetta’s achievements in biomolecular structure prediction and design over the past fifteen years. Still, the energy function is far from complete and will continue to evolve long after this publication. Thus, we hope this document will serve as an important resource for understanding the foundational physical and mathematical concepts in the energy function. Furthermore, we hope to encourage both current and future Rosetta developers and users to understand the strengths and shortcomings of the energy function as it applies to the scientific questions they are trying to answer.

## Supporting Information

Supporting Information File: `RosettaEnergyFunctionReview_Alford`_`etal_SupportingInfo`.pdf

## Author Contributions

Wrote the manuscript: RFA, JRJ, ALF, JJG

Analysis Scripts and Examples: RFA, JRJ, MSP, JJG

Expertise on protein energy terms and editing: ALF, FPD, MJO, HP, PB, MVS, RLD, BK, JJG

Expertise and description of non-protein energy terms and editing: PDR, KK, VKM, JWL, RB

## Acknowledgements

While our author list reflects main contributors to this paper, the Rosetta energy function work would not be possible without the collaboration of the entire Rosetta Commons: a community of scientists, engineers, and software developers that have worked together for almost 20 years. We thank the community for their continued dedication to empowering users to ask and answer interesting biological questions with Rosetta. We thank Sergey Lyskov for development of the benchmark server that enables continuous and transparent energy function testing. We also thank Morgan Nance and Henry Lessen for helpful comments on the manuscript.

## Footnotes

a In Rosetta, σ*i,j* has the same definition as the *r*_*i,j*_^min^ variable in CHARMM.

---

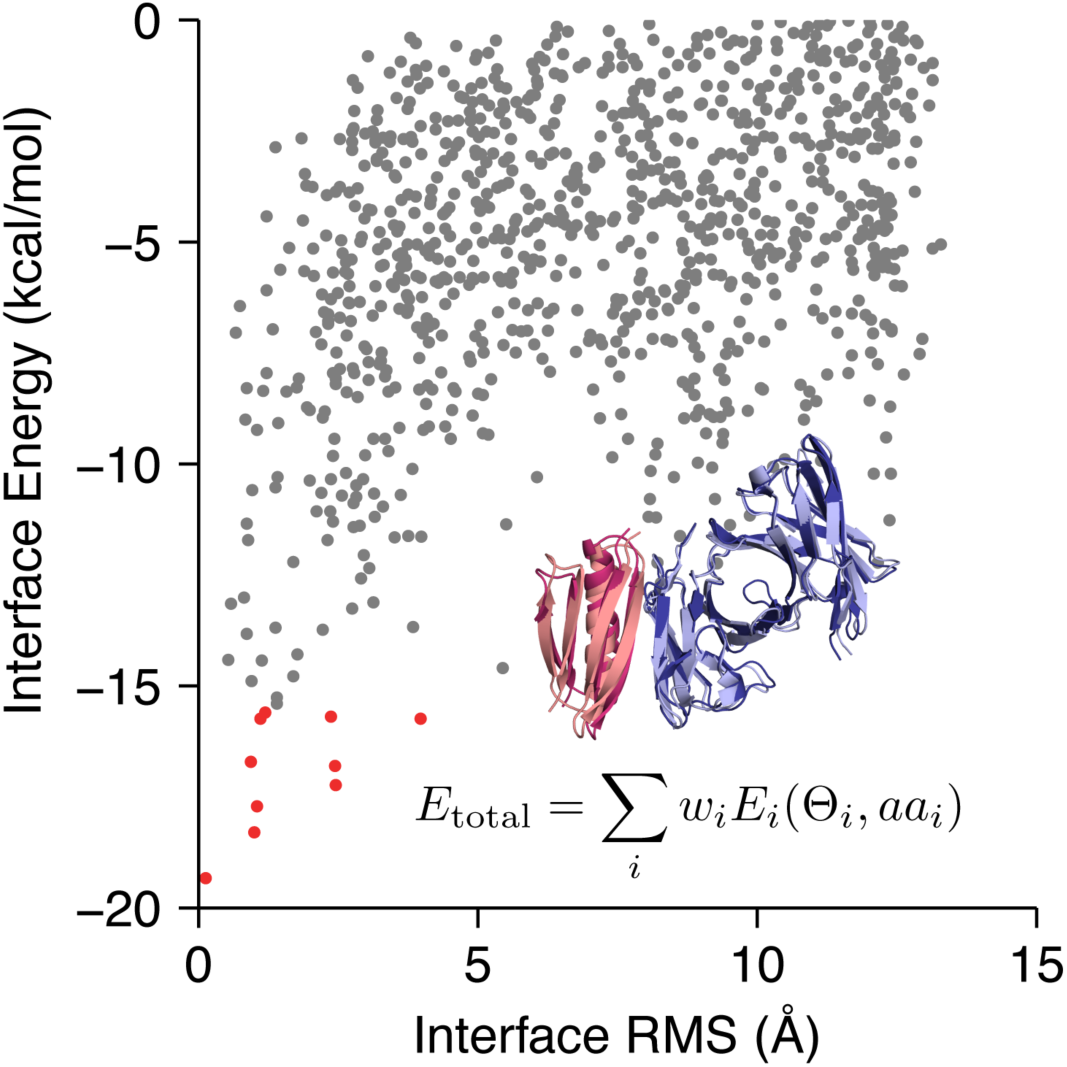
Table of Contents Graphic

